# Evolution of chlorophyll degradation is associated with plant transition to land

**DOI:** 10.1101/2021.10.07.463469

**Authors:** Isabel Schumacher, Damian Menghini, Serguei Ovinnikov, Mareike Hauenstein, Nick Fankhauser, Cyril Zipfel, Stefan Hörtensteiner, Sylvain Aubry

## Abstract

Colonization of land by green plants (Viridiplantae) some 500 million years ago was made possible by large metabolic and biochemical adaptations. Chlorophyll, the central pigment of photosynthesis, is highly photo-active. In order to mitigate deleterious effects of pigment accumulation, some plants have evolved a coordinated pathway to deal with chlorophyll degradation end-products, so-called phyllobilins. This pathway has been so far mostly unravelled in *Arabidopsis thaliana*. Here, large-scale comparative phylogenomic coupled to an innovative biochemical characterization strategy of phyllobilins allow a better understanding how such a pathway appeared in Viridiplantae. Our analysis reveals a stepwise evolution of the canonical pheophorbide *a* monooxygenase/phyllobilin pathway. It appears to have evolved gradually, first in chlorophyte’s chloroplasts, to ensure multicellularity by detoxifying chlorophyll catabolites, and in charophytes outside chloroplasts to allow adaptation of embryophytes to land. At least six out of the eight genes involved in the pathway were already present in the last common ancestor of green plants. This strongly suggests parallel evolution of distinct enzymes catalysing similar reactions in various lineages, particularly for the dephytylation step. Together, our study suggests that chlorophyll degradation accompanied the transition from water to land, and was therefore of great importance for plant diversification.

## Introduction

Photosynthetic organisms on Earth are largely dominated by green plants (Viridiplantae). They comprise two major clades: embryophytes (land plants) and algae (chlorophytes and streptophyte algae/charophytes). Embryophytes are thought to be derived from a single clade within streptophytes and diverged some 450-500 million years ago (Wickett *et al.,* 2014; de Vries & Archibald, 2018). The photosynthetic apparatus is derived from ancestral endosymbiotic cyanobacteria, which use chlorophyll as one of the main pigments in their photosystems and light-harvesting complexes (Xiong & Bauer, 2002). While our understanding of the biochemistry and evolution of chlorophyll biosynthesis is fairly comprehensive (Tanaka & Tanaka, 2006; Chew & Bryant, 2007), the pigment’s turnover and degradation have mostly been characterized in angiosperms with no actual understanding of when the pathway appeared and whether these processes are conserved in other plants (Kuai *et al.,* 2017). Despite chlorophyll degradation being one of the most visually striking biochemical reactions on Earth, with an estimated one billion tons turnover yearly (Hendry *et al.,* 1987), its occurrence outside angiosperms and the associated evolutionary advantages to degrade chlorophyll using a tightly controlled enzymatic pathway are relatively unclear.

Most of the experimental work deciphering the pathway responsible for chlorophyll degradation has been performed in angiosperms, namely on senescing leaves of *Arabidopsis thaliana* (hereafter Arabidopsis) (Hörtensteiner *et al.,* 2019) and few other ripening fruits of crop plants (Gómez *et al.,* 2013). It is referred to as the “pheophorbide *a* oxygenase (PAO)/phyllobilin pathway” (Kuai *et al.,* 2017; Hörtensteiner *et al.,* 2019;, Fig. 1A). In green leaves of Arabidopsis, chlorophyll *a* and *b* are embedded in the photosystem and light harvesting antennas. The pigment is subject to an active turnover in the so-called “chlorophyll cycle” (Tanaka & Tanaka, 2019). Upon leaf senescence, while thylakoids start to disassemble, the chlorophyll degradation pathway starts. It is generally considered to be divided into two successive phases: first, the release of the pigment from the thylakoid membrane up to the opening of the porphyrin ring, and second, the detoxification and transport of the linear tetrapyrrole until its final storage in the vacuole.

**Figure 1.**
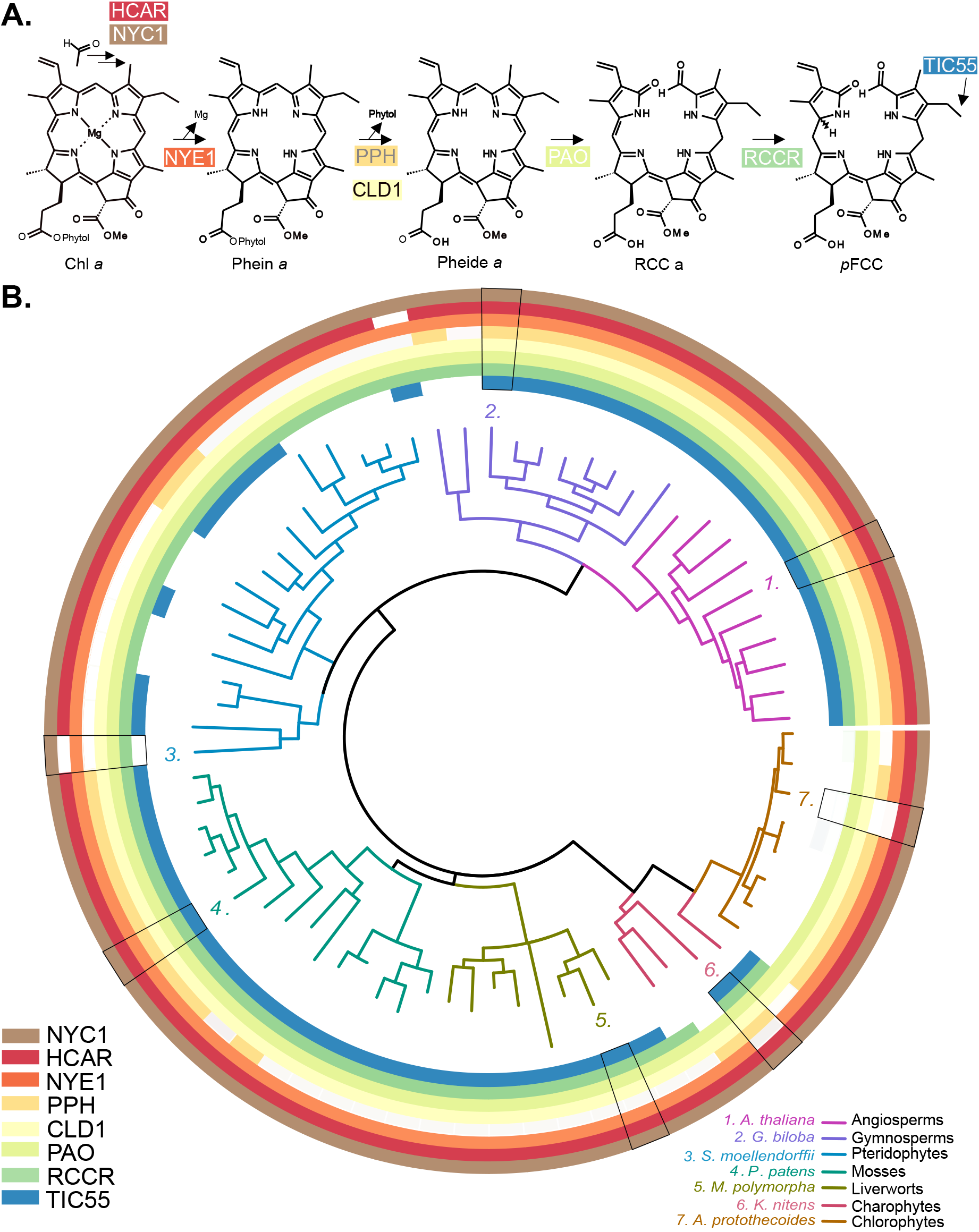
Evolution of the canonical PAO/phyllobilin chlorophyll degradation in Viridiplantae. (A) The pathway of chlorophyll degradation in Arabidopsis leaves. Only early steps of the chloroplastic portion of the PAO/phyllobilin pathway that is further studied is shown here. Further subsequent steps to the pFCC export to the chloroplast are detailed here: Kuai *et al.,* 2017. (**B) Presence of Gene Homologs Encoding Chlorophyll Degradation across 72 Plant and Algal Species.** Each concentric ring indicates an enzyme and the presence (coloured) or absence (empty) of a homolog for that particular gene. Phylogenetic tree with full species names can be find in SFigure 1. NYC1, NONYELLOW COLORING1, CHLOROPHYLL *b* REDUCTASEs; HCAR, HYDROXY-CHL *a* REDUCTASE; NYE1, NON YELLLOWING 1; PPH, PHEOPHYTINASE; PAO, PHEOPHORBIDE *a* MONO-OXYGENASE; RCCR, RCC REDUCTASE; TIC55, TRANSLOCON AT THE INNER CHLOROPLAST ENVELOPE 55; Phein a, pheophytin a; Pheide *a*, pheophorbide *a*; RCC, red chlorophyll catabolite; pFCC, primary FCC.

The first committed step to chlorophyll degradation is the conversion of chlorophyll *b* to *a* by a two-step reaction catalysed by NONYELLOW COLORING 1 (NYC1) and HYDROXYMETHYL CHLOROPHYLL a REDUCTASE (HCAR) (Kusaba *et al.,* 2007; Sato *et al.,* 2009; Horie *et al.,* 2009; Meguro *et al.,* 2011). The pigment is then subsequently processed by the Mg-dechalatase NON YELLOWING (NYE1) (Shimoda *et al.,* 2016), the dephytylase PHEOPHYTINASE (PPH) (Schelbert *et al.,* 2009), and the PHEORPHORBIDE A OXYGENASE (PAO) that catalyses the irreversible opening of the porphyrin ring (Pruzinska *et al.,* 2003). After these coordinated reactions, which leave no detectable trace of intermediaries under normal physiological condition, a first linear tetrapyrrole is produced: the red chlorophyll catabolite (RCC; Rodoni *et al.,* 1997). In Arabidopsis, it is directly converted to a primary fluorescent catabolite (pFCC) by the RED CHLOROPHYLL CATABOLITE REDUCTASE (RCCR) (Mühlecker *et al.,* 1997; Wuthrich *et al.,* 2000; Hörtensteiner *et al.,* 2000). pFCCs are then modified and further exported from the chloroplast. While the first part of the pathway appears relatively straight-forward, the fate of pFCC outside the chloroplast appears more diverse and involves various detoxifying enzymes. For example, in Arabidopsis, CYP89a9 and MES16 catalyse C1 deformylation and O8^4^ demethylation respectively (Christ *et al.,* 2012, 2013, 2016). Alone in Arabidopsis, nine distinct phyllobilins – defined as any linear tetrapyrrole that is derived from chlorophyll – have been characterized to date (Kräutler & Hörtensteiner, 2014), suggesting many more plant phyllobilin-modifying enzymes are still to be identified (Kräutler, 2016).

From a physiological point of view, maintaining a complex and costly enzymatic pathway to degrade chlorophyll in senescent leaves may allow nitrogen remobilization from both the pigment itself and their chlorophyll-binding proteins (Hortensteiner & Feller, 2002; Wang & Grimm, 2021). Indeed, about two thirds of leaf nitrogen are localized in chloroplast. However, the nature of the exact link between the pigment degradation and other senescing processes remains elusive. Importantly, leaf senescence appears to be a relatively recent invention in plants (Thomas et al., 2009), and nitrogen remobilization unlikely to be a conserved evolutionary trigger for degrading chlorophyll in all chlorophyll-containing species. Another possible reason to actively degrade chlorophyll is linked to its phototoxic properties. Some mutants impaired in the PAO/phyllobilin pathway accumulate toxic intermediates: accumulation of pheophorbide *a* or RCC strongly impact tissue integrity (Pruzinska *et al.,* 2003, 2005; Aubry *et al.,* 2020), while the end-products, mostly phyllobilins and monopyrroles, are photoinactive (Kräutler, 2016).

Our understanding of the PAO/phyllobilin pathway is therefore largely biased and limited to our knowledge of the pathway in angiosperms. For example, the vast majority of phyllobilins detected in angiosperms appears to be structurally related (with opening of the porphyrin ring at the “northern” C1 position); therefore, suggesting that the PAO/phyllobilin pathway of degradation is largely conserved (Hörtensteiner *et al.,* 2019). Some recent data showing presence of phyllobilins (*iso*-phillobilanones) in ferns (*Pteridium aquilinum*) suggest that the PAO/phyllobilin pathway is present outside angiosperms (Erhart *et al.,* 2018). Indeed, chlorophyll catabolites were identified in the green alga *Auxenochlorella protothecoides* (Gossauer, 1994). Chlorophyll degradation might even have evolved outside the green lineage, in marine protists like the dinoflagellates *Pyrocystis lunula* (Wu *et al.,* 2003). In these two particular cases, the exact nature of the enzymes involved remains to be determined. In *P. lunula*, the opening of the ring that produces luciferin occurs at a different position of the porphyrin ring as compared to plant PAO products (Wu *et al.,* 2003). This is likely due to an independent evolution of the chlorophyll degradation pathway in this clade. On the contrary, *A. protothecoides* accumulates RCC-like compounds when grown heterotrophically. It excretes RCCs into the surrounding media, which therefore strongly suggests a PAO-like activity in this alga (Hortensteiner *et al.,* 2000).

Taking advantage of recent advances in phylogenomics; especially, the increased number of high-quality genomes and transcriptomes available, we performed a comparative analysis of homologous genes involved in the chlorophyll degradation in the green lineage (Cannell *et al.,* 2020; Harris *et al.,* 2020). Orthologs are genes that show some homology and diverged after a species split rather than after duplication (Fitch, 1970; Emms & Kelly, 2015). We merged and computed genomes and transcriptomes from a subset of 72 species representative of seven major lineages in plant evolution (SFig. 1). We based our analysis of potential homologs of the known chlorophyll degradation Arabidopsis proteins. In addition, we performed targeted metabolomics to detect of the end-products of this pathway (phyllobilins) in some representative species of these lineages.

We aimed here to determine when during plant evolution a dedicated pathway to degrade chlorophyll might have evolved. It remains unclear whether a unique strategy (*i*.*e*. the PAO/phyllobilin pathway) for chlorophyll catabolites detoxification has been recruited. Eventually, we seek to understand the physiological relevance of detoxifying chlorophyll and identify families or lineages where potential further studies could shed more light on the chlorophyll degradation pathway’s evolutionary trajectory.

## Results

### Identification of chlorophyll catabolic enzymes in plants by ortholog’s inference

Despite some pioneering work in barley and fescue, the vast majority of the biochemistry of the PAO/phyllobilin pathway has been characterized in Arabidopsis. Recent works highlight the potential of orthology inference, using tools like Orthofinder (Emms & Kelly, 2020), to better understand evolution of particular traits like stomata development (Harris *et al.,* 2020) or various specialized metabolic pathways (Cannell *et al.,* 2020). Here, using a similar approach, we performed orthology inference on 72 land plants and algae that have their genomes sequenced and represent each of the major lineages in plant evolution: angiosperms, gymnosperms, lycophytes, bryophytes (mosses and liverworts), charophytes and chlorophytes (Fig. 1B). Out of these species, 87.3 % of genes could be assigned to 24,479 orthogroups. Then, orthogroups containing potential orthologs for eight major enzymes involved in the PAO/phyllobilin pathway were identified. Focusing on each enzymatic step at a time, we then checked for presence/absence of orthologs in Viridiplantae and inferred the pathway evolution in these species (Fig. 1B).

### The “chlorophyll cycle” enzymes are well conserved throughout the green lineage

Conversion of chlorophyll *a* to *b* and its reverse reaction is catalysed by a set of four enzymes working in the so-called “chlorophyll cycle”. Chlorophyll *a* is converted into chlorophyll *b* by the CHLOROPHYLL *A* OXYGENASE (CAO), that is often also considered the “last” step of chlorophyll synthesis. Given that chlorophyll *b* is also found in cyanobacteria (Satoh *et al.,* 2001), it was expected that all the genes responsible for its synthesis would be present in all eukaryotic photosynthetic organisms. The vast majority of genes involved in chlorophyll synthesis, including *CAO*, are indeed present in all 72 species analysed here (SFig. 2).

The chlorophyll *b* to *a* conversion is performed in a two-step process: NYC1 reduces chlorophyll *b* to 7-hydroxymethyl chlorophyll and HCAR subsequently converts this intermediate into chlorophyll *a* (Zhao *et al.,* 2020). Here, we focused on the chlorophyll *b* to *a* conversion (*i*.*e*. NYC1 and HCAR), as strictly speaking the two necessary steps that trigger chlorophyll degradation (Shimoda *et al.,* 2012). Indeed, PAO and NYE1 do not accept “*b*” pigments as substrates (Shimoda *et al.,* 2016). Our analysis identifies orthologs of these two enzymes throughout the entire Viridiplantae (Fig. 1B, Fig. S3). Interestingly, a paralog (*i*.*e*. a gene that derived from the same ancestral gene by duplication) of NYC1, NYC-LIKE 1 (NOL), belongs to the same orthogroup but is phylogenetically clearly distinct from NYC1: it is absent from chlorophytes, suggesting NOL might have originated from a relatively recent gene duplication event (SFig. 3).

### Evolution of the Mg-dechelation

The *STAYGREEN* gene (*SGR*, also referred to as *NYE1*) was one of the first genetic loci indicative of a pathway responsible for chlorophyll degradation (Armstead *et al.,* 2006, 2007; Aubry *et al.,* 2008). Its role as a Mg-dechelatase has only been shown recently (Shimoda *et al.,* 2016). *NYE1* genes were identified in all Viridiplantae lineages (Fig. 1B). Intriguingly, only two species did not show *NYE1* orthologs: the charophyte *Klebsormidium nitens* and the chlorophyte *A. protothecoides*. Given the presence of phyllobilins in *A. protothecoides* (SFig. 4), and the fact that *NYE1* is present in all other green algae studied (both charophytes and chlorophytes), these absences might be likely due to uneven genome quality/assembly and not to gene loss (Cannell *et al.,* 2020). In Arabidopsis, three paralogues (*NYE1, SGR2* and *SGL*) are involved in the dechelating reaction and might differ in their substrate specificity (Sakuraba *et al.,* 2014; Lin *et al.,* 2016). Phylogenetic analysis suggests they all derived from a recent gene duplication event (Fig. 2A). Most species analysed here show more than one copy of the *NYE1* gene (for example, four copies in *G. biloba*, Fig. 2A). While the enzymatic steps preceding and succeeding the Mg-dechelation are apparently being encoded by single copy genes, the biochemical reason for having multiple copies of this particular gene remains unclear but might be related in some cases to the maintenance of photosystem homoeostasis.

**Figure 2.**
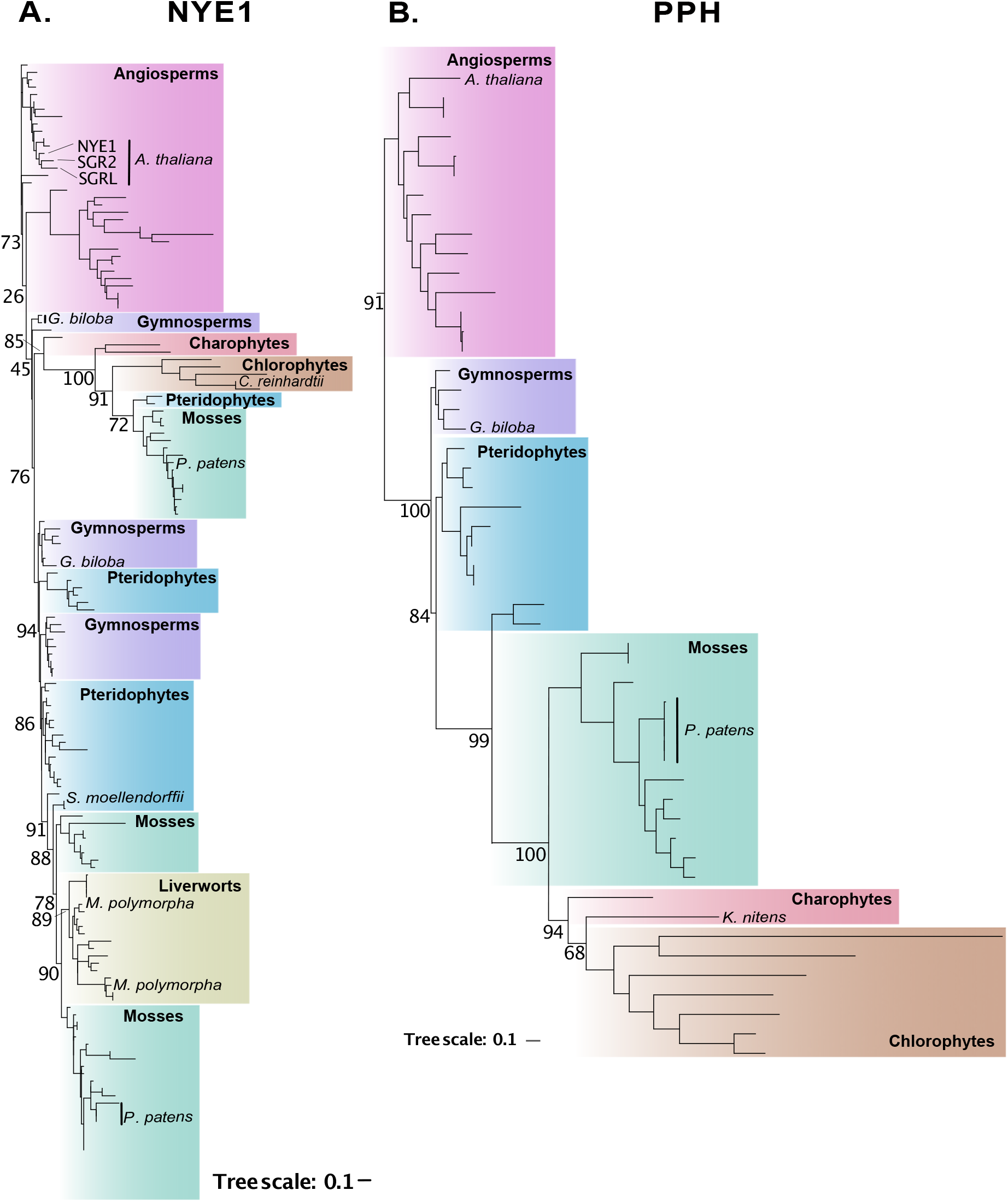
Evolution of NYE1 and PPH in the Viridiplantae (A) Maximum likelihood phylogenetic tree of NYE1. In Arabidopsis, NYE1 paralogs SGR2 and SGRL appear to be the result of relatively recent duplications events. Interestingly, the chlorophyte NYE1 ortholog in *C. reinhardtii*, shown to be involved in chlorophyll dephytylation appears to be clustering within a branch containing mosses, gymnosperms, charopytes and pteridophytes. A distinct branch from liverworts till gymnosperms appears to be related to NYE1 but without known function **(B) Maximum likelihood phylogenetic tree of PPH.** PPH appears to be derived from a common ancestor of land plants and the gene was apparently loss in liverworts as well as some genera in pteridophytes and mosses. Only bootstrap support values for key branches are shown beside each corresponding branch.

### Multiple enzymes might have been recruited for dephytylation reactions during evolution

The subsequent step to the Mg-dechelation in Arabidopsis is the removal of the phytol chain from pheophytin, which is catalysed by PHEOPHYTINASE (PPH) (Schelbert *et al.,* 2009). Removal of the phytol chain is a key step that allows the release of the pigment from the apoproteins. Impairments in PPH induces a stay-green phenotype in several angiosperms like Arabidopsis (Schelbert *et al.,* 2009), *Lolium perenne* (Zhang *et al.,* 2016) and pear (*Pyrus communis*) (Cheng & Guan, 2014). Despite being functionally conserved in all angiosperms studied, our analysis shows that *PPH* is not uniformly present among all Viridiplantae. There is no *PPH* homolog in pteridophytes, lycophytes, hornworts and liverworts (Fig. 1B). Interestingly, some streptophyte algae and chlorophytes apparently do contain *PPH* homologs (Fig. 2B).

Since the lycophyte *S. moellendorffii* and the liverwort *M. polymorpha* are both able to produce phyllobilins (Table 1 and SFig. 6), it is therefore possible that these species express a distinct dechelating enzyme catalysing the conversion of pheophytin *a* into pheophorbide *a*. The hydrolysis of the phytol chain can be catalysed by esterases and lipases, which are relatively large families of enzymes in plant genomes (Schelbert *et al.,* 2009). Forty-two chloroplast-localized α/β hydrolases were screened during PPH identification in Arabidopsis (Schelbert *et al.,* 2009). At least two other enzymes had previously been described to dephytylate chlorophyll: chlorophyllase (CLH, Schenk *et al.,* 2007; Hu *et al.,* 2015) and CHLOROPHYLL DEPHYTYLASE 1 (CLD1, (Lin *et al.,* 2016). CLH has been first thought to dephytylate chlorophyll but its subcellular localization is still matter of debates and may depend on leaf age (Schenk *et al.,* 2007; Tian *et al.,* 2021). CLH has also been shown to be involved in phyllobilin release upon defence activation against chewing herbivores (Hu *et al.,* 2015). In this model, CLH accesses its substrate only after destruction of the chloroplastic membranes. CLH is therefore an unlikely candidate for dephytylation in the PPH-free lineages we identified here. Another dephytylase, CLD1, that is localized to the chloroplast, has been shown to be involved in chlorophyll steady-state turnover and possible recycling of the pigment in Arabidopsis (Lin et al., 2016). CLD1 is an interesting candidate for performing dephytylation in lycophytes, pteridophytes, liverworts and hornworts (SFig. 5); although, this requires functional confirmation. Having at least three enzymes (CLH, PPH, CLD1) being able to catalyse the chlorophyll dephytylation suggests that there is a wider range of possible hydrolases that may catalyse this reaction outside angiosperms. More work is required to identify the enzyme(s) responsible for dephytylation in these lineages.

**Table 1.**
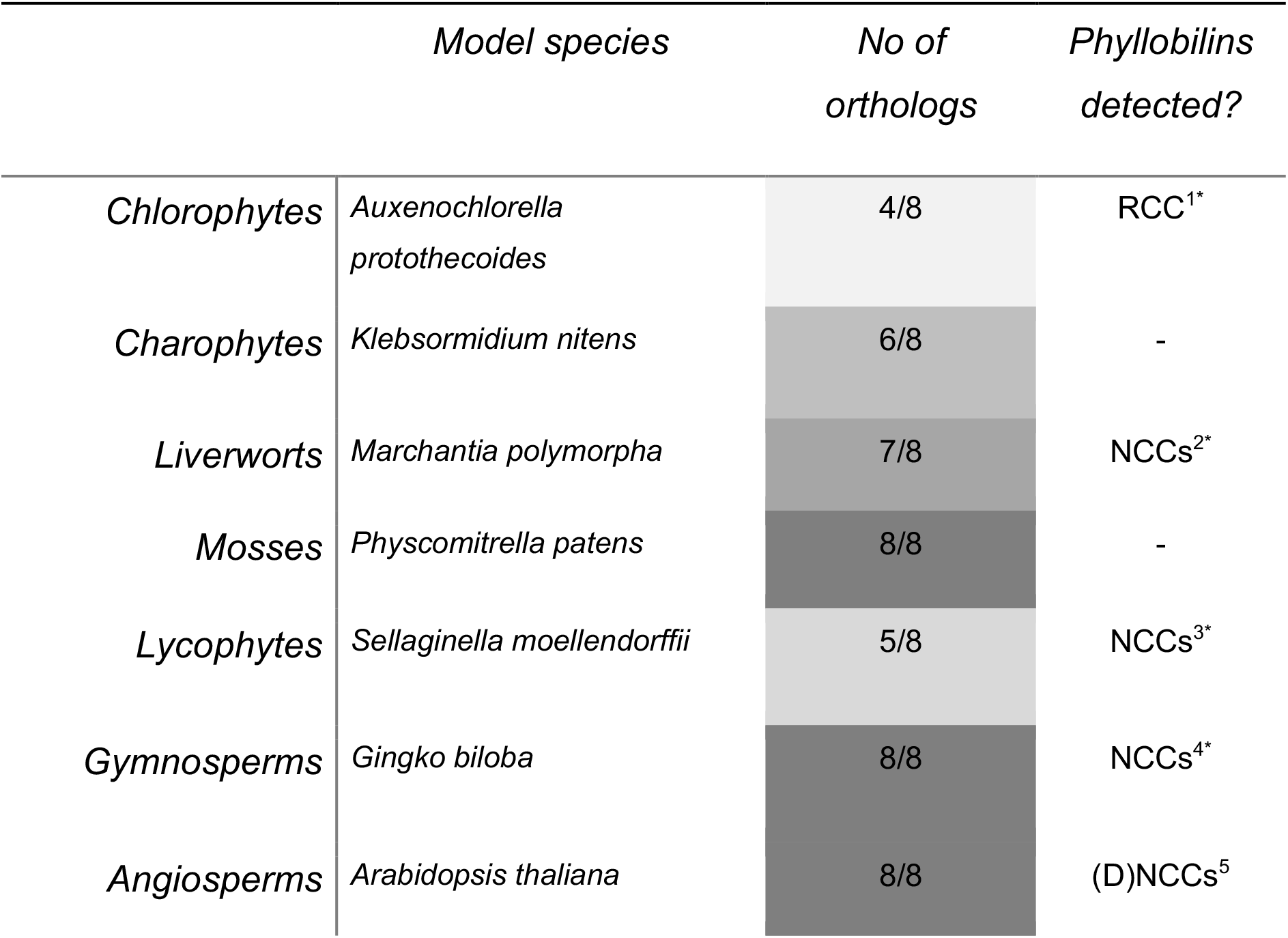
Detection of homologs of the PAO pathway and phyllobilins across the green clade suggest some possible alternative pathways to chlorophyll degradation. *This study. ^1^*Ap*RCC1 and *Ap*RCC2 (SFig. 3) ^2^previously undescribed NCC with multiple hydroxylation (SFig. 3), ^3^ & ^4^NCC similar to DNCC_618 (Kuai *et al.,* 2017), ^5^see Christ et al., 2016. RCC, red chlorophyll catabolite; NCC, colourless chlorophyll catabolite.

|  | *Model species* | *No of orthologs* | *Phyllobilins detected?* |
| --- | --- | --- | --- |
| *Chlorophytes* | *Auxenochlorella protothecoides* | 4/8 | RCC<sup>1*</sup> |
| *Charophytes* | *Klebsormidium nitens* | 6/8 | - |
| *Liverworts* | *Marchantia polymorpha* | 7/8 | NCCs<sup>2*</sup> |
| *Mosses* | *Physcomitrella patens* | 8/8 | - |
| *Lycophytes* | *Sellaginella moellendorffii* | 5/8 | NCCs<sup>3*</sup> |
| *Gymnosperms* | *Ginkgo biloba* | 8/8 | NCCs<sup>4*</sup> |
| *Angiosperms* | *Arabidopsis thaliana* | 8/8 | (D)NCCs<sup>5</sup> |

### Evolution of the irreversible opening of the porphyrin ring

One of the major steps of the chlorophyll degradation pathway is the opening of the porphyrin ring catalysed by PAO (Pruzinska *et al.,* 2003). PAO converts pheophorbide *a* into a red chlorophyll catabolite (RCC). PAO is a Rieske-type mononuclear non-heme iron oxygenase that has five close-relatives in Arabidopsis: PAO, CHLOROPHYLL *A* OXYGENASE (CAO), CHOLINE MONO-OXYGENASE (CMO), PROTOCHLOROPHYLLIDE-DEPENDENT TRANSLOCON COMPONENT 52 (PTC52) and TRANSLOCON AT THE INNER CHLOROPLAST ENVELOPE 55 (TIC55) (Gray et al., 2004). Interestingly, PTC52, which has been suggested to have a flavone hydroxylase activity in basil (*Ocimum basilicum*) (Berim *et al.,* 2014), belongs to the same orthogroup as PAO but not TIC55 (Fig. 3A). PAO orthologs are present across the green lineage (Fig. 1B). Potential *PAO* orthologs in chlorophytes group in a distinct “PAO-like” clade that also encompasses some charophytes, mosses and liverworts sequences (Fig. 3A). Alignment of the Rieske-type and mononuclear iron domain of the *A. protothecoides* PAO show only about 40% identity to other PAO orthologs (SFig. 7). Despite some early indications of a monooxygenase involvement in this reaction by application of specific inhibitors (Hortensteiner *et al.,* 2000), the actual ability of this gene to catalyse pheophorbide *a* production remains to be experimentally confirmed.

**Figure 3.**
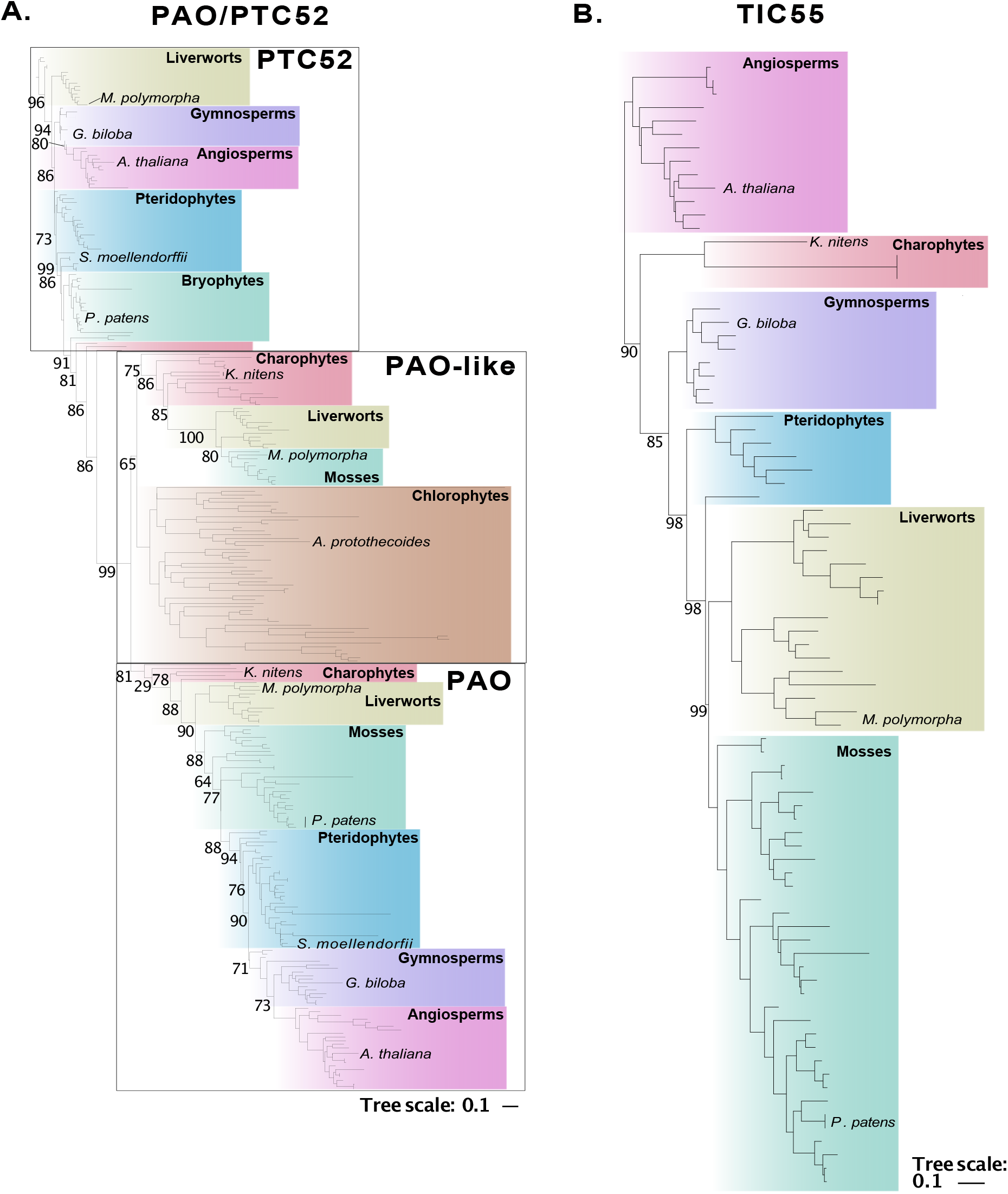
Evolution of PAO and TIC55 and PPH in the Viridiplantae. (A) Maximum likelihood phylogenetic tree of PAO. Three genes appear to be clustered in one orthogroups that might have been present in the most common ancestor of plants: PTC52, PAO and PAO-like. While a *bona fide* PAO appears to be present in charophytes, the relevance of distinction of a PAO-like subfamily will remain to be studied further. Noteworthy, *A. protothecoides* ortholog do belongs to the PAO-like subfamily. **(B) Maximum likelihood phylogenetic tree of TIC55.** TIC55 does appear as an innovation of the charophytes and remain monophyletic with few gene duplication (expect in mosses). TIC55 being a Rieske-type monooxygenase, it might have derived a pre-existing enzyme, while relationship to the PAO or CAO ancestors is unclear. Only bootstrap support values for key branches are shown beside each corresponding branch.

TIC55 localizes in the chloroplast in Arabidopsis and catalyses the C3^2^ hydroxylation of pFCC (Hauenstein *et al.,* 2016). Multiple hydroxylated phyllobilins have been detected (Christ & Hörtensteiner, 2014; Kräutler, 2016), but their physiological relevance during leaf senescence remains unclear. TIC55 is absent from chlorophytes and is only present in some charophytes. It also appears missing in most pteridophytes (Fig. 1B and Fig. 3B). While hydroxylation of phyllobilin modifies the polarity of phyllobilins to be exported, TIC55’s presence is not mandatory for a functional degradation of phyllobilins and therefore not a very significant marker of the pathway’s evolution.

### Evolution of enzymatic steps leading to the phyllobilin’s export from the chloroplast

During leaf senescence, it is essential that photoreactive chlorophyll intermediates do not accumulate, in order to avoid toxicity and tissue damages. Therefore, photoreactive RCCs produced by PAO need to be detoxified quickly. RCCR reduces the C15/16 double bond of RCC to produce less photoreactive primary fluorescent catabolites (pFCC) (Pruzinska *et al.,* 2005). The conversion from a closed ring into a linear tetrapyrrole is possibly the trigger allowing export of the catabolites from the chloroplast to the cytosol (by an yet unknown transporter) (Kuai *et al.,* 2017). RCCR orthologs are conserved in streptophytes (Fig. 1B and Fig. 4), while it appears absent from chlorophytes. This result is consistent with reported RCCR activities from liverworts, mosses and gymnosperms (Pruzinska *et al.,* 2007). In a similar line, the green algae *A. protothecoides* and *Parachlorella kessleri* excrete large amounts of RCC in their growing medium when grown under low nitrogen and heterotrophic conditions (their chemical structure is described in SFig. 6; Gossauer, 1994; Hortensteiner *et al.,* 2000). While *A. protothecoides* and *P. kessleri* have evolved a dedicated detoxification of RCC, the extent to which a similar strategy is employed by other chlorophytes remains unclear.

**Figure 4.**
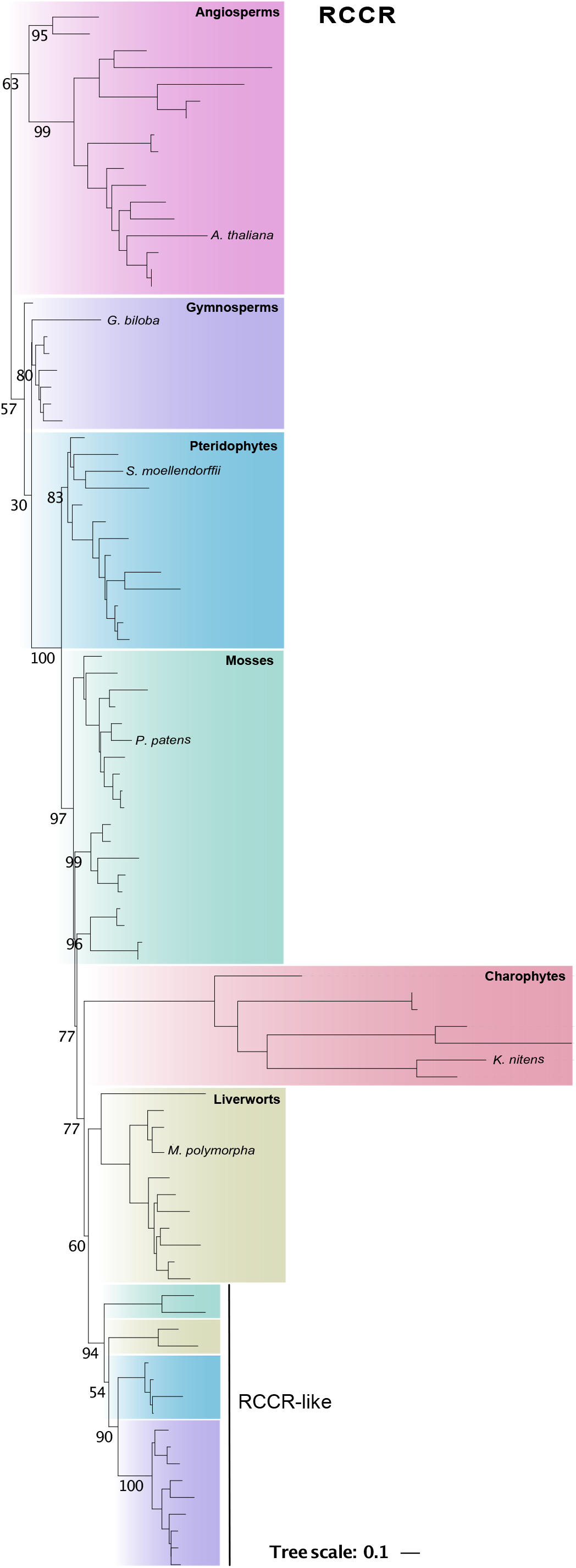
Phylogenetic tree of RCCR. RCCR appears as an innovation from the charophytes and is present in all other clades. Interestingly, there is a distinct sub-family of RCCR-like genes in gymnosperms, mosses and pteridophytes of unknown function. Only bootstrap support values for key branches are shown beside each corresponding branch.

### Detection of phyllobilins in key species is consistent with ortholog’s inference

We then focused on key species for each of the seven lineages for experimentally confirming the presence of the pathway’s end-product, and logically infer the conservation of the PAO/phyllobilins pathways across the green lineage (Table 1). The presence/absence of phyllobilin was tested first using dark-induced senescence in detached leaves (or chlorophyll-containing tissues) for the embryophytes Arabidopsis, *S. moellendorffii, G. biloba, P. patens* and *M. polymorpha* (see Materials and Methods). Degradation of chlorophyll was induced in the two algae *A. protothecoides* and *K. nitens* after switching into heterotrophic conditions (low nitrogen and darkness). Phyllobilins could be detected in five of the seven clades investigated (Table 1). Despite several attempts and a diverse range of media and conditions tested (variations in nitrogen content before and after the dark incubation, change in carbon sources and duration of the growth and dark incubation, bubbling vs. mixed cultures, etc…), no phyllobilins could be identified neither from *K. nitens* nor *P. patens* (data not shown). However, shift to heterotrophic growth of *A. protothecoides* induced excretion of two types of RCC in the culture media (SFig. 6). *Ap-*RCC1 and *Ap-*RCC2 structures only differ by a single carboxymethyl group bound to the isocyclic ring of chlorophyll in C8^2^ position (SFig. 6). In Arabidopsis, this reaction is catalysed by MES16 (Christ et al., 2012), suggesting that MES16-like activities are present in chlorophytes. Unfortunately, MES16 belongs to an orthogroup gathering many *α/β*-hydrolases, making it very hard to properly identify orthologs. *M. polymorpha* incubated in the dark accumulates multiple phyllobilins (SFig. 6). At least four distinct compounds could be identified, including *Mp-*NCC-4, for which MS-MS fragmentation pattern was shown to be identical to NCC-662 (Kuai *et al.,* 2017; Hörtensteiner *et al.,* 2019). It seems that multiple successive rounds of hydroxylation are added to the core *Mp-*NCC4 in *M. polymorpha*, but their exact position on the porphyrin ring remains to be determined (SFig. 6). The presence of NCCs in *M. polymorpha* confirms an active PAO/phyllobilin pathway in liverworts, as predicted by our phylogenomic approach (Fig. 1B). But these results also highlight the complexity and versatility of cytosolic side-chain modification.

### Large scale phyllobilin’s detection show the extent of structural diversity

In order to better characterise the diversity of chlorophyll catabolites in phyllobilin-producing species, we took advantage of our laboratory localization, at the heart of the Botanical Garden of the University of Zürich. During three consecutive autumns, we harvested naturally senescing leaves and extracted phyllobilins out of 203 species and analysed them by LC-MS/MS. Phyllobilins were detected in 183 species (90 %) (STable 2).

To increase our chance to identify phyllobilins, we screened the measured masses against an *in silico* produced list of ‘diagnostic ions’ with their predicted masses (Figure 5B). We based ourselves on one of the simplest structures of naturally occurring phyllobilin, a C1 deformylated pyroRCC (m/z 556.2686, Figure 5). We then predicted all possible combinations of masses that could be originating from the 12 modifications over 8 positions (Fig. 5). These modifications are namely C1 deformylation, C3 hydroxylation, malonylation/glucosylation or combination of both, C2-C4 hydroxymethylation, C2 semi-reduction, C8 demethylation, C8 demethylesteration, C12 hypermodification, C18 dihydroxylation, rearrangement of the carbon backbone where the C5/C6 is shifted to a C5/C7 bond (Hörtensteiner et al., 2019). A total of 4,992 hypothetical structures could be listed, in which all 26 known phyllobilins could be retrieved from, as well as 15 profiles that are potentially yet undescribed phyllobilins (STable 2 and Fig. 5C). Interestingly, major structural modifications like the conversion in *iso*-phyllobilanones (m/z 619.2762 [M+H]+ and m/z 635.2712 [M+H]+), recently reported in *P. aquilinum* (Erhart et al., 2018), could also be detected in most ferns and gymnosperms studied here (STable 2). Taken together, the diversity of cytosolic phyllobilins observed here could appear surprising considering the high degree of conservation of the (preceding) chloroplastic steps and might be indicative of a more general strategy of cells to deal with potentially toxic catabolites.

**Figure 5.**
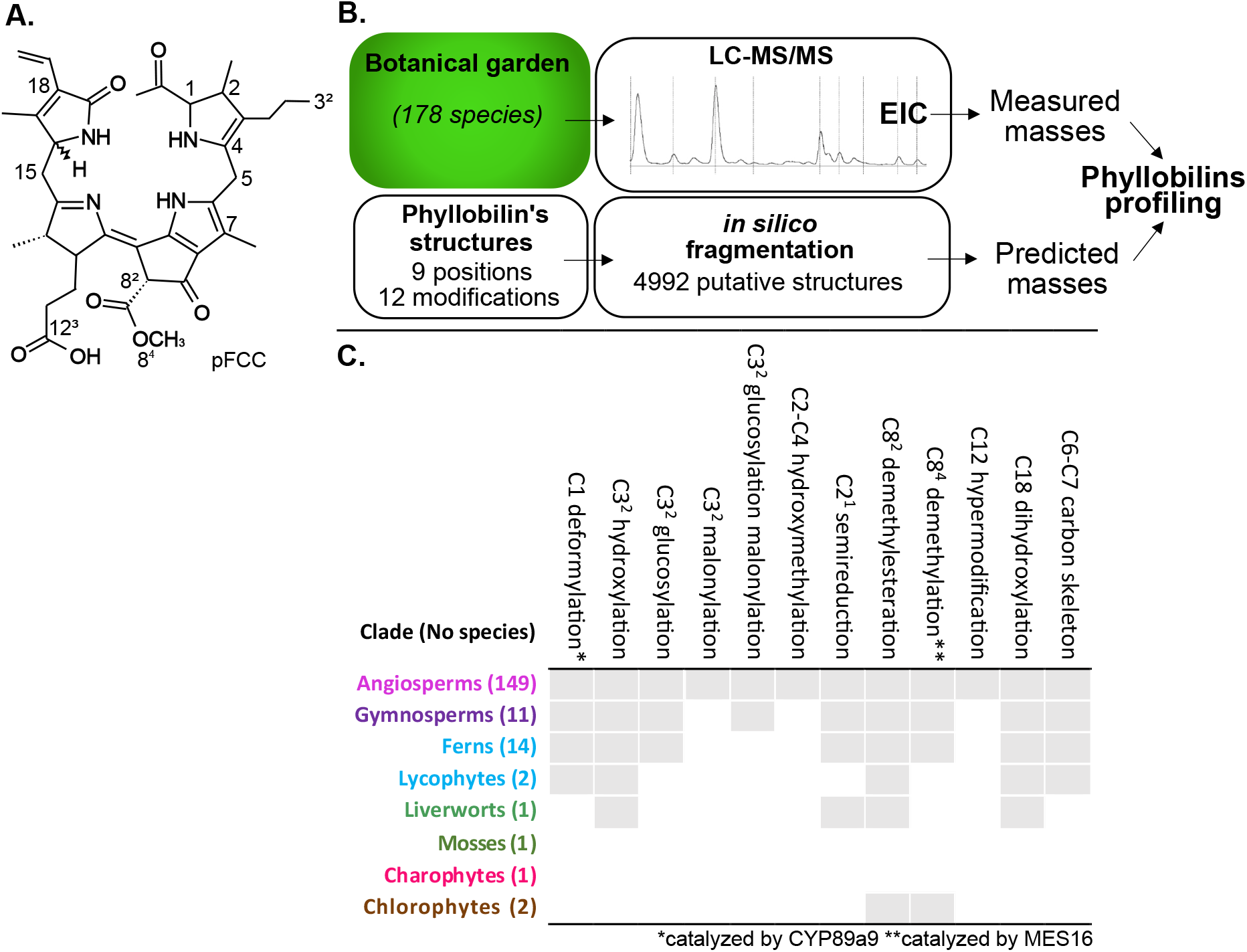
Diversity of cytosolic phyllobilins. **(A)** Mapping of the various positions of modifications based on primary FCC. Modifications give rise to fluorescent phyllobilins (DFCCs and FCCS) that are isomerized further in the cytosol. **(B)** Strategy used for identification of de novo phyllobilins. **(C)** Table summarizing all phyllobilin’s side chain modification described in this study.

## Discussion

Taking advantage of our knowledge of the PAO/phyllobilin pathway in angiosperms and the extensive genomic resources covering Viridiplantae, we decipher here the evolution of the chlorophyll degradation in this clade. This approach was complemented by detection of the pathway’s end-products, phyllobilins. Our analysis provides a comprehensive view over possible ways by which plants have dealt with phototoxic chlorophyll catabolites during evolution.

### Chloroplastic degradation of chlorophyll may have facilitated transition to multicellularity

Some unicellular chlorophytes like *A. protothecoides* (SFig. 4) and *P. kessleri* (Gossauer, 1994) degrade chlorophyll in their chloroplast, most likely using a PAO-like protein (Fig. 1B & 2A), process phyllobilins to a certain extent (presence of modified RCCs in *A. protothecoides*, Fig. S4), and eventually excrete phyllobilins into the media (Hortensteiner *et al.,* 2000). It remains unclear whether the ability to produce RCC in *A. protothecoides* can be generalized to other chlorophytes, or is a *de novo* adaptation of this particular species, which allows the switch between autotrophic and heterotrophic growth without accumulation of phototoxic compounds (Gao *et al.,* 2014). Our data suggest that, in chlorophytes, the enzymatic machinery is present but the actual degradation of chlorophyll (and associated gene expression) most likely depends on the lifestyle of each alga.

Interestingly, genes recruited in the PAO/phyllobilin pathway appear to have an ancestral and often distinct function and substrates. For example, an ortholog of *NYE1* in *Chlamydomonas reinhardtii* (belonging to the NYE1-like orthogroup, Fig. 2A), has been proposed to facilitate photosystem II biosynthesis instead of acting as a magnesium dechelatase in plants (Chen *et al.,* 2019). Homologs of *NYE1* could be identified in some glaucophytes and non-photosynthetic bacteria, but neither in cyanobacteria nor rhodophytes (Obata *et al.,* 2019). This might be indicative of a recruitment of *NYE* in plants via horizontal gene transfer succeeding the cyanobacteria primary endosymbiosis. Meanwhile, the large variation of substrate specificity and dechelating efficiency of the homologs may question their physiological relevance. In other words, the extent to which the -*in vitro*- ability of a protein to dechelate magnesium is a good proxy for testing its actual orthology to the angiosperm’s NYE/SGR remains unclear. A similar cyanobacterial origin for *HCAR* has also been suggested: it may have derived from a cyanobacterial Ferredoxin-dependent divinyl reductase (F-DVR), responsible for the reduction of an 8-*vinyl* group in the chlorophyll biosynthesis (Ito & Tanaka, 2014). Therefore, the presence of *HCAR, NYC1* and *NOL* across all species from the green lineage may not be surprising.

PAO activity, as a mono-oxygenase, depends obviously on molecular oxygen presence; therefore, production of RCC must have logically succeeded the appearance of oxygenic phototrophs, in very comparable way haem is degraded in bilins (Zhang *et al.,* 2021). *PAO* and *TIC55* also appear to have homologs in several oxygenic cyanobacteria (Gray et al., 2004). As the ability to convert chlorophyll *a* to *b* by CAO has been suggested to appear several times in the prokaryotic prochlorophytes (Tomitani *et al.,* 1999), it is therefore possible that *PAO* and *TIC55* did originate from duplication events from *CAO*. Chl *b* conversion by CAO is thought have participated to aerobic photosynthesis evolution (Tomitani *et al.,* 1999). But again, ancestral monooxygenase homologs are very difficult to distinguish without proof of their biochemical activity.

In charophytes, despite no phyllobilins being detected in *K. nitens* (some pheophorbide *a* trace could be detected in *Chara vulgaris*; STable 2), all genes encoding proteins converting chlorophyll to RCCs are present (Fig. 1B). This suggests that charophytes may degrade chlorophyll, although the growth conditions triggering chlorophyll degradation in these species remain elusive. While phototrophic lineages, including charophyte algae, successfully established on land multiple times independently (Lewis & McCourt, 2004), the land plant macroflora originates from a unique streptophyte ancestor (de Vries & Archibald, 2018). Comparative genomic supports two genetic innovation bursts during emergence of multicellularity in streptophytes and terrestrialization (embryophytes) (Bowles *et al.,* 2020).

Taken together, our results suggest that the chloroplastic portion of the PAO/phyllobilin pathway (*i*.*e*. from chlorophyll up to RCC) was already present in the common ancestor of land plants (Fig. 6). The recruitment of the PAO-RCCR couple allowed a swift conversion of the highly phototoxic pheophorbide *a* and its subsequent export (as a linear RCC or pFCC) from the chloroplasts (Hirashima *et al.,* 2009; Aubry *et al.,* 2020). This appeared as a key innovation crucial for allowing multicellularity of phototropic organisms by dealing with highly phototoxic chlorophyll catabolites embedded in thylakoid membranes.

**Figure 6.**
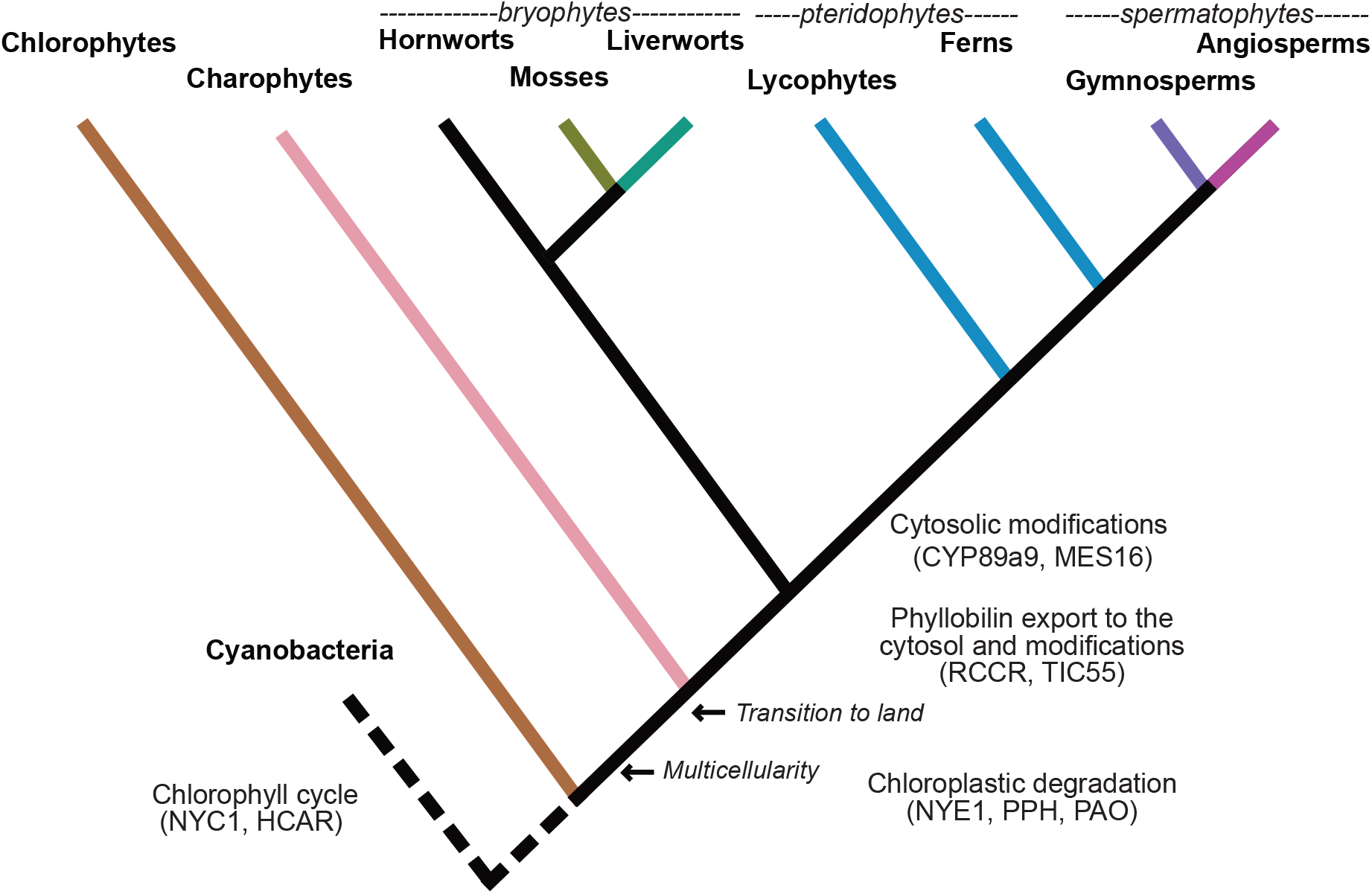
A model for the evolution of the PAO/phyllobilin pathway in plants. Cladogram of the major plant lineages and evolution of key traits that marked the transition from aquatic to a terrestrial life.

### Phyllobilin production evolved coincidentally to water-to-land transition

Given the phototoxicity of chlorophyll catabolites intermediates, especially for multicellular organisms where excretion is not an option, it is interesting to observe that the enzymatic steps that take place within the chloroplasts appear under a much higher selective pressure compared to cytosolic processes. There is in fact only two catabolites (pFCC and hydroxy-pFCC) known to be exported from the chloroplasts, that are subsequently converted in more than 40 different phyllobilins in streptophytes (STable 2). This diversity of cytosolic activities is most likely due to the high redundancy of detoxifying processes that appear to ensure an extensive range of enzymatic activities (reductases, hydrolases…) and of relatively high promiscuity. Low substrate specificity allows the enzymes to react with virtually any toxic/xenobiotic compound that may be detrimental to the cellular activity. Interestingly, hydroxylations of specialised metabolites (diterpenoids) that also involve cytochrome P450s were shown to be involved in plant herbivore’s defense, while preserving plants from their own toxicity (Li *et al.,* 2021). A similar function for phyllobilin’s accumulation remains to be investigated (Kräutler, 2016). It is therefore possible that these cytosolic steps allowed to mitigate the impact of light stress associated to terrestrial lifestyle (and partly associated to the chlorophyll catabolites phototoxicity) and were central to the transition to land.

### Lower genomic resolution for the cytosolic phyllobilin-modifying steps

Despite at least nine different phyllobilins detected in senescing Arabidopsis leaves (Hörtensteiner *et al.,* 2019), only three enzymes have been associated with phyllobilin side-chain modifications, namely TIC55, CYP89a9 and MES16 (Christ *et al.,* 2012, 2013; Hauenstein *et al.,* 2016). CYP89a9 belongs to the large and highly interrelated multi-gene family of cytochrome P450 oxidase and MES16 to the *α/β*-hydrolase family (Christ *et al.,* 2012, 2013). It is therefore not surprising that they both clustered in two large orthogroups, which makes the identification of proper candidate genes difficult. For example, the cytochrome P450 family comprises 245 members in Arabidopsis and has undergone rapid expansion and neofunctionalization (Nelson & Werck-Reichhart, 2011; Du *et al.,* 2016). Therefore, screening for intra-specific (natural) variations between phyllobilins (*e*.*g*. deformylated, demethylated or conjugated phyllobilins) that could be related to some specific loci by genome-wide association study might be a complementary strategy to identify more cytosolic enzymes involved in chlorophyll degradation.

### Diversity of the cytosolic modifications across phyllobilin-producing species

Monitoring phyllobilins in 183 species via a ‘Botanical Garden metabolomics’ approach provided a unique view on the diversity of cytosolic modifications originating from one single pigment: chlorophyll. While the sampling is clearly biased towards angiosperms, it provides a unique perspective on the structural diversity of phyllobilins. Surprisingly, about 10 % of the samples did not contain any detectable phyllobilins. This could be due to their respective leaf phenologies: despite using an extensive harvesting period, not all species or organs do induce senescence and degrade chlorophyll at the same time. Particularly mosses and charophytes were hardly degrading chlorophyll despite numerous attempts to induce senescence artificially in these species. Moreover, some species like tobacco do not accumulate phyllobilins to detectable amounts, while degrading chlorophyll. This might be due to a particularly quick transfer to acidic vacuoles or possibly to large amounts of secondary metabolites that might limit phyllobilin’s extraction. While our innovative strategy to predict *in silico* and screen any possible sorts of phyllobilin improved the pool of molecules to be screened for, it is still possible that some catabolites were simply missed during the analysis.

Since their first chemical characterization in 1996 in barley, a constantly increasing number of phyllobilins have been described (Kräutler, 2016; Christ *et al.,* 2016). Two major families of phyllobilins can be distinguished: type I that are non-fluorescent (NCCs), 1-formyl, 19-oxobilins and type II, dioxobilin-type non-fluorescent (DNCCs) (Kräutler, 2014, 2016). Within these two categories, a wide range of side-chain modifications have been reported. Some modifications, like the C_3^2^_ hydroxylation (Fig. 5C) are largely shared, while many others phyllobilins are restricted to only few species. For example, hypermodified FCC with functionalization in C^12^ position have been detected in banana (*Musa acuminata*) and *Spathiphyllum wallisii* (Moser *et al.,* 2012; Vergeiner *et al.,* 2015). In *Ulmus glabra*, a bicycloglysidic NCC connecting C_12^3^_ and C_3^2^_ was reported (Scherl *et al.,* 2016). Some other modifications appear restricted to some clades, like the major rearrangement of the carbon backbone that has been initially observed in *P. aquilinum* but found here to be happening in ferns and gymnosperms (Erhart et al., 2018, STable 2).

Finally, using this large dataset, no particular phylogenetic pattern could be observed that would indicate a common ancestor to any of the various modifications downstream of TIC55 (STable 2). Phyllobilins accumulating in senescent leaves, while having sometimes similar structures, may in many instances result from distinct (promiscuous) enzymatic activities. In terms of physiological relevance, this indicates that a relatively diverse and promiscuous set of cytosolic detoxication reactions have been recruited to change phyllobilin’s polarity towards vacuolar export. But in contrast to the highly conserved chloroplastic portion of the pathway, these reactions are likely shared with other detoxication processes and do not necessarily evolve in a coordinated manner. More extensive analyses in other clades will be indicative of the actual extent of complexity of the cytosolic detoxifying processes during land plant diversification.

## Conclusions

Combining homology inference and biochemical characterization over a wide range of species covering the green lineage has proven to be a promising approach to help understanding evolution of biological pathways. Completion of more and better-quality genomes (particularly for large and repetitive genomes like in gymnosperms) will surely improve the quality of ortholog’s identification at the species level (Cannell *et al.,* 2020). Noteworthy, most gymnosperms that do not degrade chlorophyll in a coordinated way (with the notable exception of *Larix decidua*) still have the entire set of PAO/Chlorophyll genes in their genome. This emphasizes the crucial role of transcriptional regulation in the onset of chlorophyll degradation. More generally, it also raises interesting questions about the possible reasons and dynamics for such genes to be kept or lost during evolution.

We could observe here a high degree of conservation of the genes involved in the early chloroplastic step of the PAO/phyllobilin pathway. Taking in consideration the few indications of potential prokaryotic homologs (at least for *HCAR, NYE1, PAO* (Gray *et al.,* 2004; Ito & Tanaka, 2014; Obata *et al.,* 2019), this raises interesting questions on their actual function in prochlorophytes and the actual process and timing by which a tightly coordinated pathway has evolved in plants. When considering chlorophyll degradation, it is also important to consider the various cellular and tissue-specific gene expression patterns that relates to changing contexts in which chlorophyll is degraded in the green lineages. Indeed, the extent of co-evolution of senescence and the hormonal cues that regulate the PAO/phyllobilin pathway remain to be determined (Thomas *et al.,* 2009; Bowles *et al.,* 2020; Lai *et al.,* 2020). In Arabidopsis, all eight genes involved in the core PAO/phyllobilins pathway are under a complex transcriptional control that involve multiple families of transcription factors that are in turn under hormonal control (Kuai *et al.,* 2017; Aubry *et al.,* 2020). Building on this work that focuses on key species, and helped by previous attempts in Arabidopsis senescent leaves (Hickman *et al.,* 2017), it would be important to identify the gene regulatory networks that control the PAO/phyllobilin pathway in all these different clades. Again here, focusing on model species with high quality genomes will be necessary to identify potential *cis* elements and transcription factors involved. It remains unclear how this pathway could be maintained under similar regulation (or at least in response to similar environmental cues, in a similar way) during the complex evolution/expansion/alteration of various transcription factor families in the green clade (Lai *et al.,* 2020). More work is needed to identify the extent of conservation of the gene regulatory networks and their connexion to various environmental/hormonal cues (Fürst-Jansen *et al.,* 2020) that regulate the PAO/phyllobilin pathway, and that degrades chlorophyll in a safe and efficient manner. Chlorophyll degradation is an overlooked yet apparently crucial aspect of the transition from water to land, and therefore is of great importance for our understanding of plant diversification.

## Material and Methods

### Evolutionary bioinformatics and phylogenetic analysis

This study used genomes and transcriptomes sequences from 72 species, all sources and accession numbers of which are summarized in STable 1. Most angiosperm genome sequences were download from Phytozome (Goodstein *et al.,* 2012), the 1KP Project (Matasci *et al.,* 2014), Fernbase (Li *et al.,* 2018), ConGenie (Sundell *et al.,* 2015) and UniProt (Bateman *et al.,* 2021)or from individual publications. Genes for the 72 selected species were sorted into orthogroups using OrthoFinder v2.5.2 software (Emms & Kelly, 2015, 2020). Orthogroups are: “collections of orthologous and paralogous genes descending from one gene in the common ancestor of all species included in the analysis” (Emms & Kelly, 2015). Among the 24,479 orthogroups created, we focused our analysis on the orthogroups that contain known Arabidopsis genes of interests.

After manual curation, gene trees were inferred for each gene following published methods (Harris *et al.,* 2020). Briefly, homologous amino acid sequences were aligned using MAFFT v7.475 (Katoh & Standley, 2013) and the resulting multiple alignment was trimmed using BGME v4.0 with a BLOSUM30 matrix (Criscuolo & Gribaldo, 2010). The clean alignment was then processed by IQ-Tree v1.6.12 using bootstrapped maximum likelihood phylogenies with an empirical profile mixture models (C10-C60) (Nguyen *et al.,* 2015). The trees were visualized and edited in iTOL v5 (Letunic & Bork, 2019).

### Plant material growth and chlorophyll degradation induction

Arabidopsis and *S. moellendorffii* were grown on soil in 16 hours day /8 hours night conditions, 100 μmol/m^2^/s, 60 % humidity. *G. biloba* leaves were harvested mid-August in the Botanical Garden of the University of Zürich. Detached leaves from these three species were dark incubated for 7 days to induce phyllobilin production. *P. patens* was grown on solid Knopp medium (Frank *et al.,* 2005) in long-day conditions and later transferred in nitrogen-free medium for dark incubation. *M. polymorpha* was grown in the same conditions as Arabidopsis (see above) on ½ Gamborg medium plates supplemented with 0.5 % sucrose. Detached thalli were dark incubated for 10 days on wet filter paper before phyllobilins were extracted. *K. nitens* was grown following a published protocol (Hori *et al.,* 2014) and nitrogen and light depletion experiments were carried out (data not shown). *A. protothecoides* (previously called *Chlorella protothecoides*) was first grown autotrophically for 10 days and then transferred to nitrogen-depleted medium for 5 days following a published protocol (Hortensteiner *et al.,* 2000).

### Phyllobilins extraction and LC-MS/MS analysis

After dark incubation and first sign of senescence, the plant materials were immediately frozen in liquid nitrogen and stored at -80 °C until further use. Prior to extraction and following a published protocol (Christ et al., 2016), the frozen samples were grind in liquid nitrogen using mortar and pestle and resuspended in five volumes (w/v) of pre-cooled extraction buffer (80 % MeOH, 20 % water and 0.1 % formic acid) (Christ *et al.,* 2016). Samples were centrifuged and supernatants were separated two times and transferred on the LC-MS system. The samples were then analysed using a LC-MS/MS Thermo Scientific Dionex Ultimate 3000 Rapid Separation LC system (Thermo Fisher Scientific, Reinach, Switzerland) equipped with a photodiode array detector coupled to a Bruker Compact ESI-QTOF (Bruker Daltonics). Phyllobilins were separated on a C18 column (ACQUITY UPLC BEH, 1.7 μm, 2.1 × 9 × 150 mm; Waters, Milford, MA USA), using a similar protocol as published previously (Christ et al., 2016). Briefly, a gradient of acetonitrile with 0.1 % [v/v] formic acid was performed with water with 0.1 % [v/v] formic acid as follows (all v/v): 30 % acetonitrile for 0.5 min, 30-70 % in 7.5 min, 70-100 % in 0.1 min and 100 % for 4 min. The instrument was set to acquire over the m/z range 50-1300, with an acquisition rate of four spectra/second. All data were recalibrated internally using pre-run injection of sodium formate (10 mM sodium hydroxide in 0.2 % formic acid, 49.8 % water, 50 % isopropanol [v/v/v]). Each sample was run in data-dependent MS/MS mode.

### Phyllobilin’s mass- and fragmentation prediction and detection

Most of the phyllobilins known so far were identified by LC-MS/MS and based on a set of diagnostic ions that are most likely fragmenting from a porphyrin ring. In order to mine for new variations in phyllobilins, LC-MS/MS data were compared to a list of *in silico* generated masses that gather all possible combinations of 12 reported modifications on 8 positions over the porphyrin’s ring (Fig. 5). A total of 4,992 hypothetical structures and their corresponding predicted masses were produced and compared to the measured masses from the MS/MS data for each species using a home-made R script (available at https://github.com/phyllomenous/phyllobilin_permutations).

## Supporting information

STable 1

STable 2

## Declarations

### Author’s contribution

IS, SH and SA designed the research, IS, MH, NF, SO, DM helped collecting data and analysing, all authors helped for the interpretation, IS, DM and SA wrote the paper.

## Acknowledgements

The authors would like to dedicate this article to our colleague, mentor and friend, the late Stefan Hörtensteiner, whose seminal work on chlorophyll degradation inspired the early steps of this work. We would like to thank Bastien Christ, Bernhard Kräutler and Philipp Schlueter for helpful comments on the manuscript. This work was supported by the Swiss National Science Foundation (#31003A_172977) and the University of Zurich.

## Supplementary Data

**Supplementary Table 1**. List of species used in this study and the data sources.

**Supplementary Table 2**. List of phyllobilins from 178 species sorted by modifications observed and sorted by position on the porphyrin ring and their corresponding phyllobilins. The phyllobilin nomenclature used here initially described in Hörtensteiner et al., 2019.

**Supplementary Figure 1.**
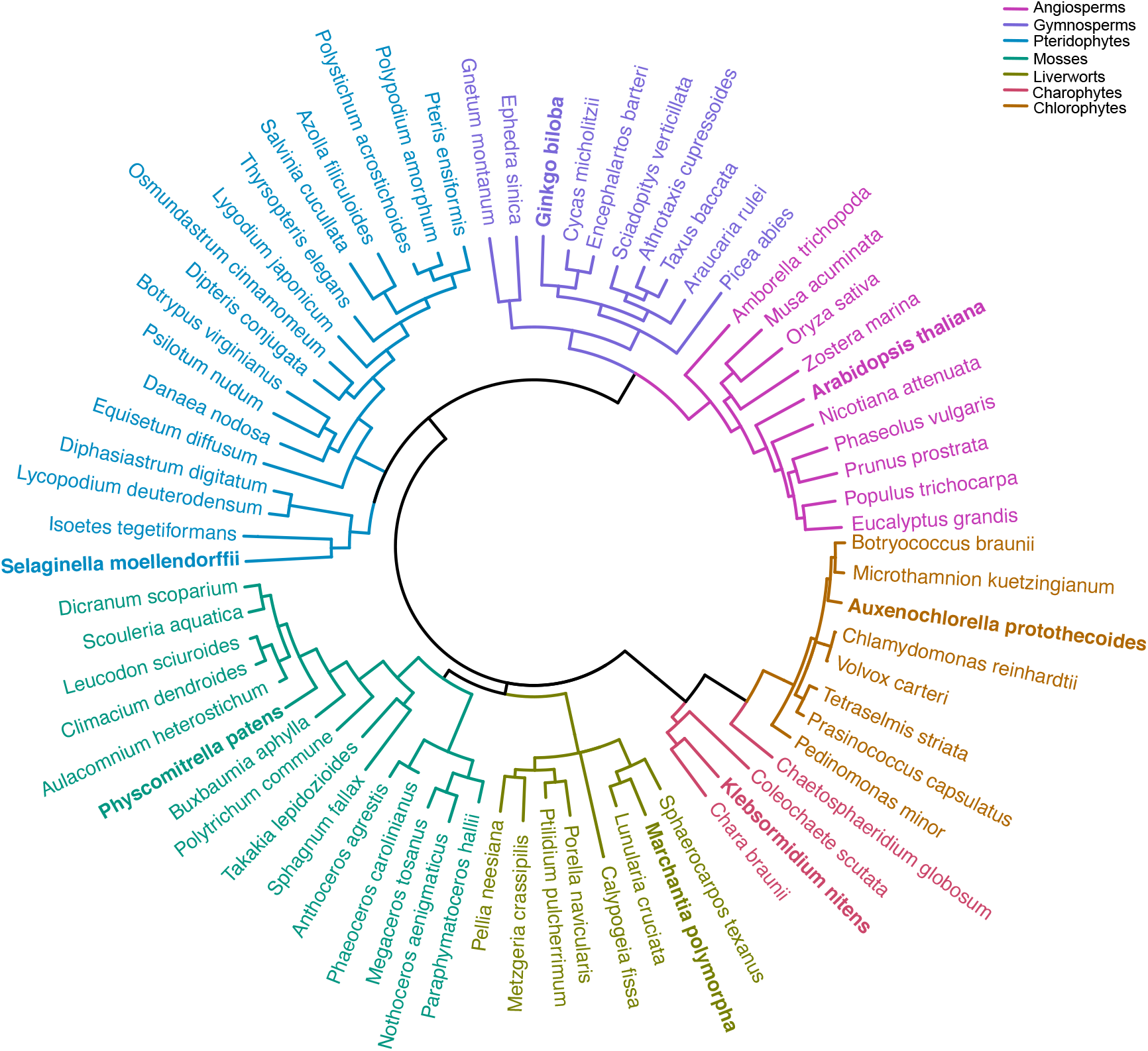
Phylogenetic tree showing evolutionary relationship between all 72 species involved in the orthogroups analysis. The same species tree was used to draw the presence/absence map in Fig. 1B. Model species used for phyllobilin detection are shown in bold.

**Supplementary Figure 2.**
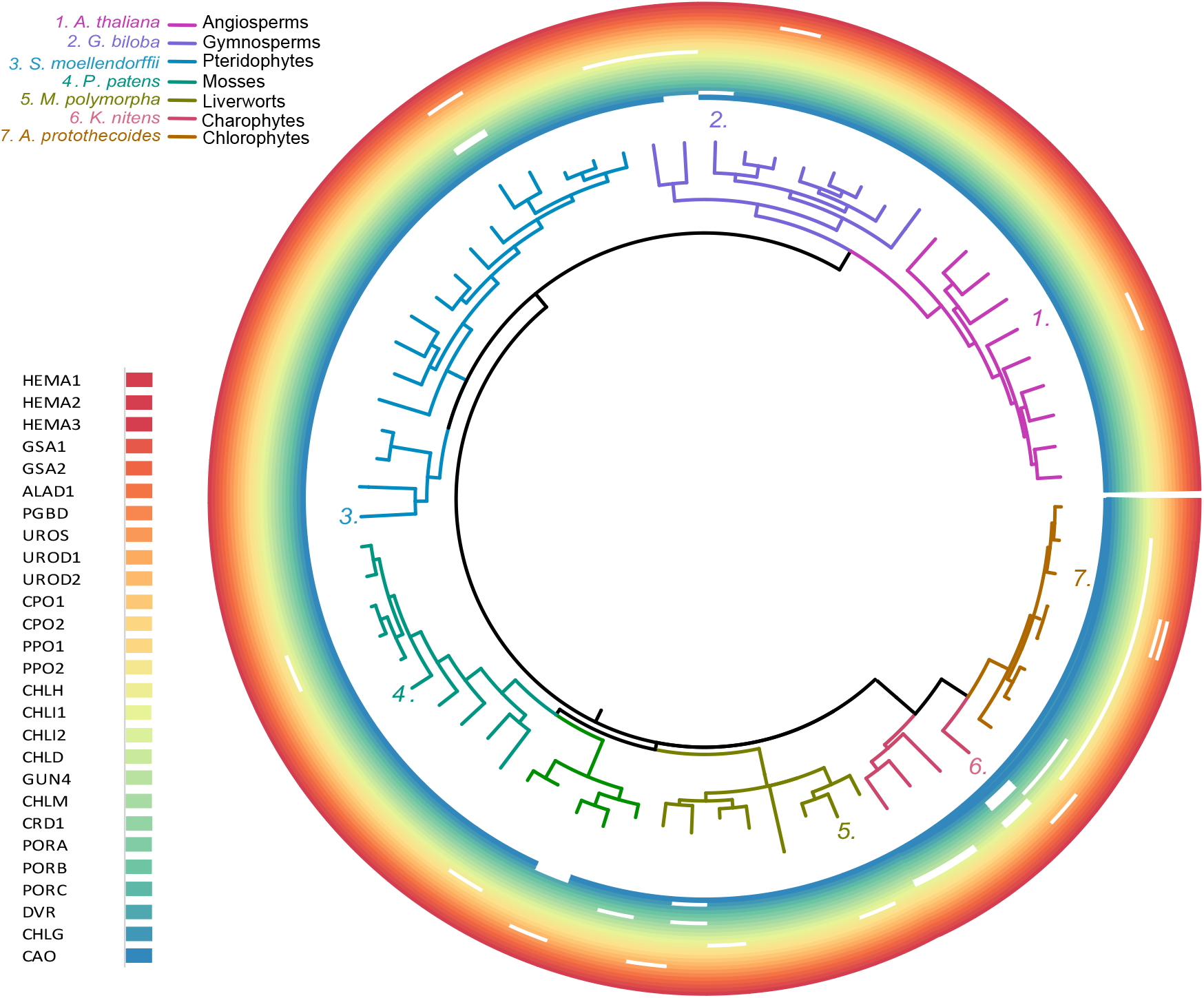
Presence of Gene Homologs Encoding Chlorophyll Synthesis across 72 Plant and Algal Species. Each concentric ring indicates an enzyme and the presence (coloured) or absence (empty) of a homolog for that particular gene. Phylogenetic tree with full species names can be find in SFigure 1. The chlorophyll synthesis pathway appears mostly conserved throughout the entire Viridiplantae.

**Supplementary Figure 3.**
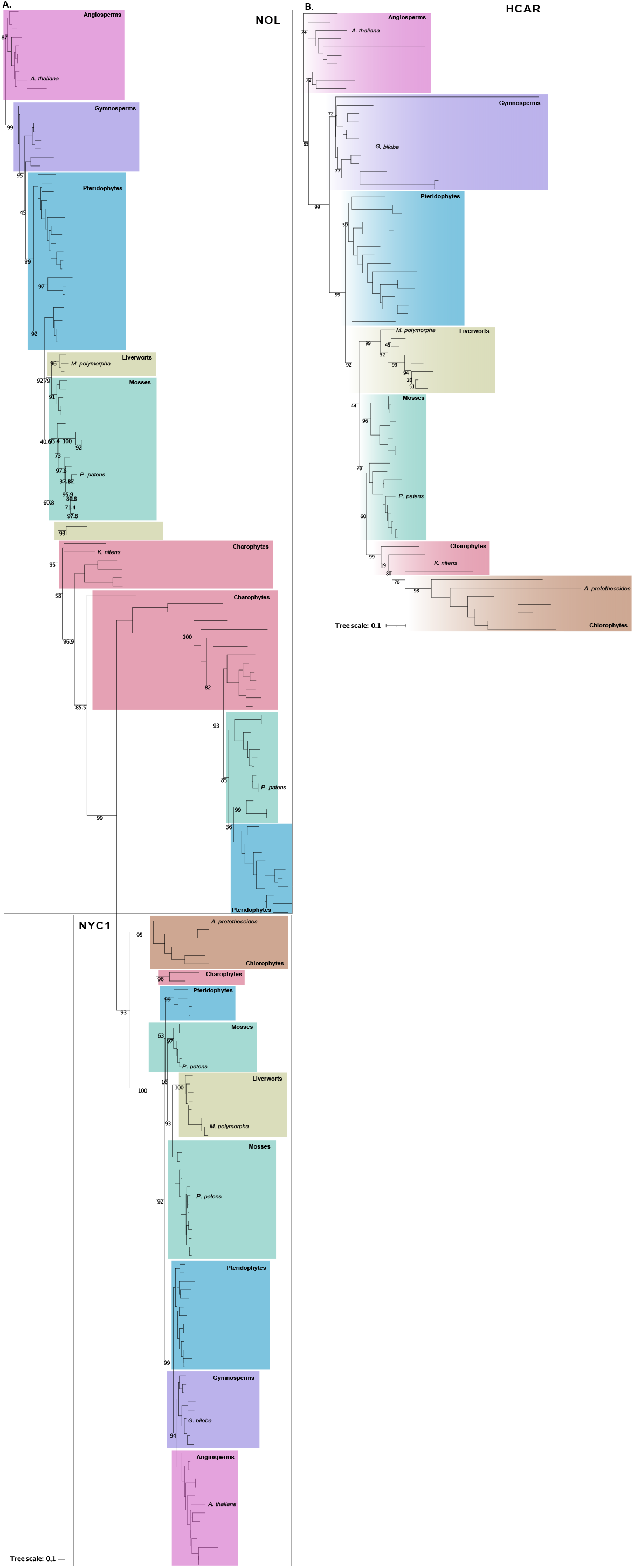
Maximum likelihood phylogenetic tree of **(A)** NYC1/NOL and **(B)** HCAR orthologs. Only bootstrap support values for key branches are shown beside each corresponding branch.

**Supplementary Figure 4.**
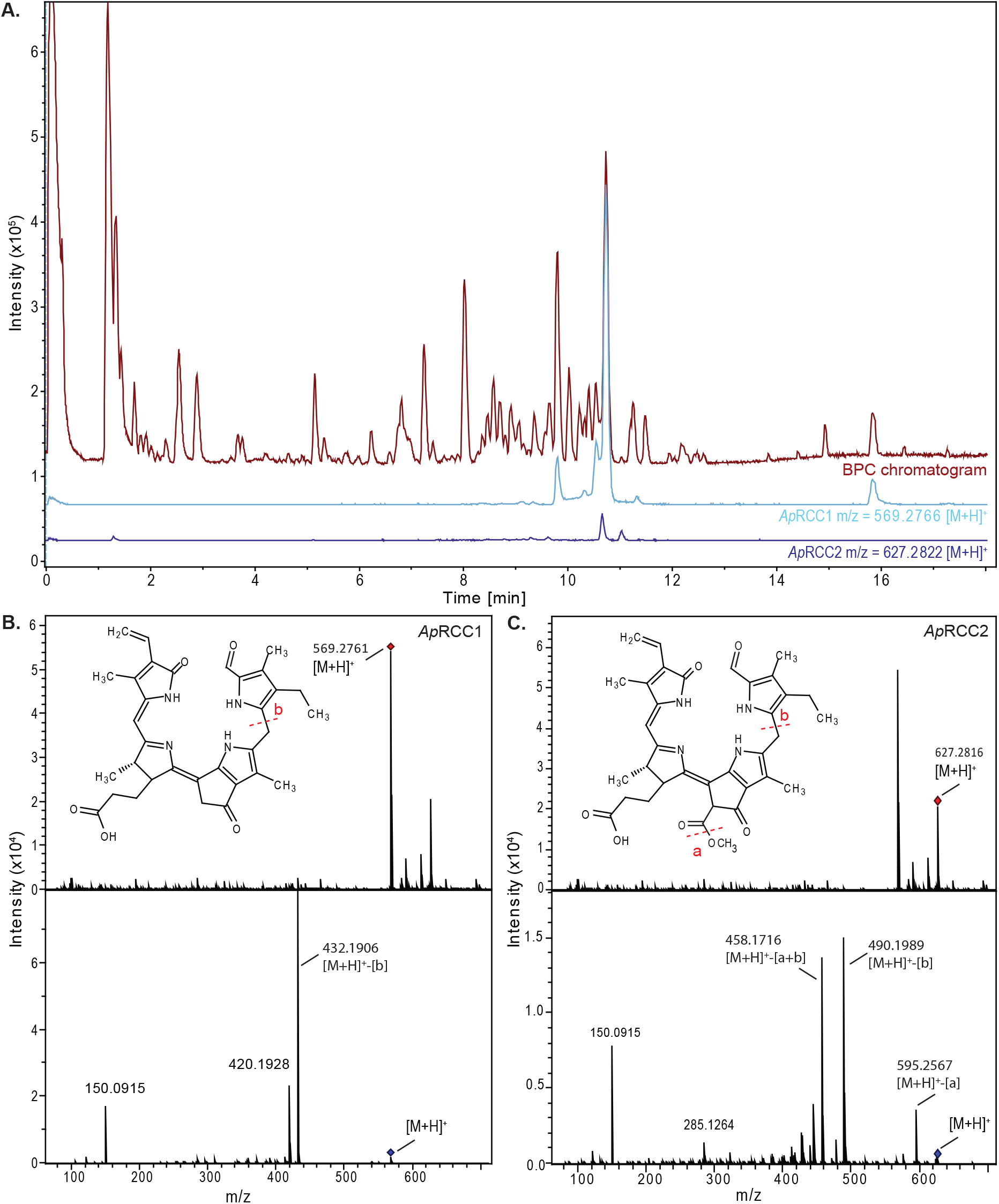
***Auxenochlorella protothecoides* excretes two types of RCC** in the growing media under heterotrophic conditions. **(A)** Base peak chromatogram BPC) of the extract and extracted ion chromatogram for ApRCC1 (light blue) and ApRCC2 (dark blue) **(B)** MS (top) and MS/MS (bottom) spectra and predicted structure models of ApRCC1, **(C)** MS (top) and MS/MS (bottom) spectra and predicted chemical structure of ApRCC2. RCC, Red Chlorophyll Catabolites.

**Supplementary Figure 5.**
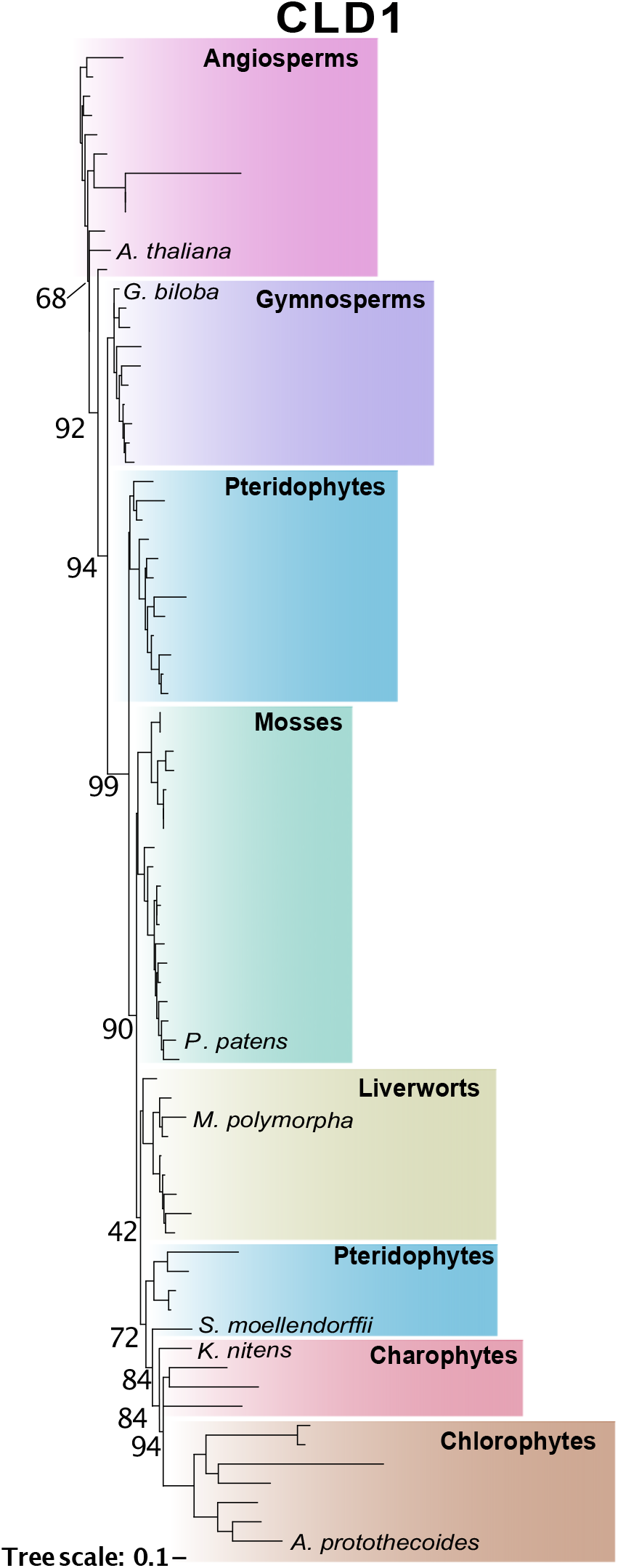
Maximum likelihood phylogenetic tree of CLD1 orthologs. Only bootstrap support values for key branches are shown beside each corresponding branch.

**Supplementary Figure 6.**
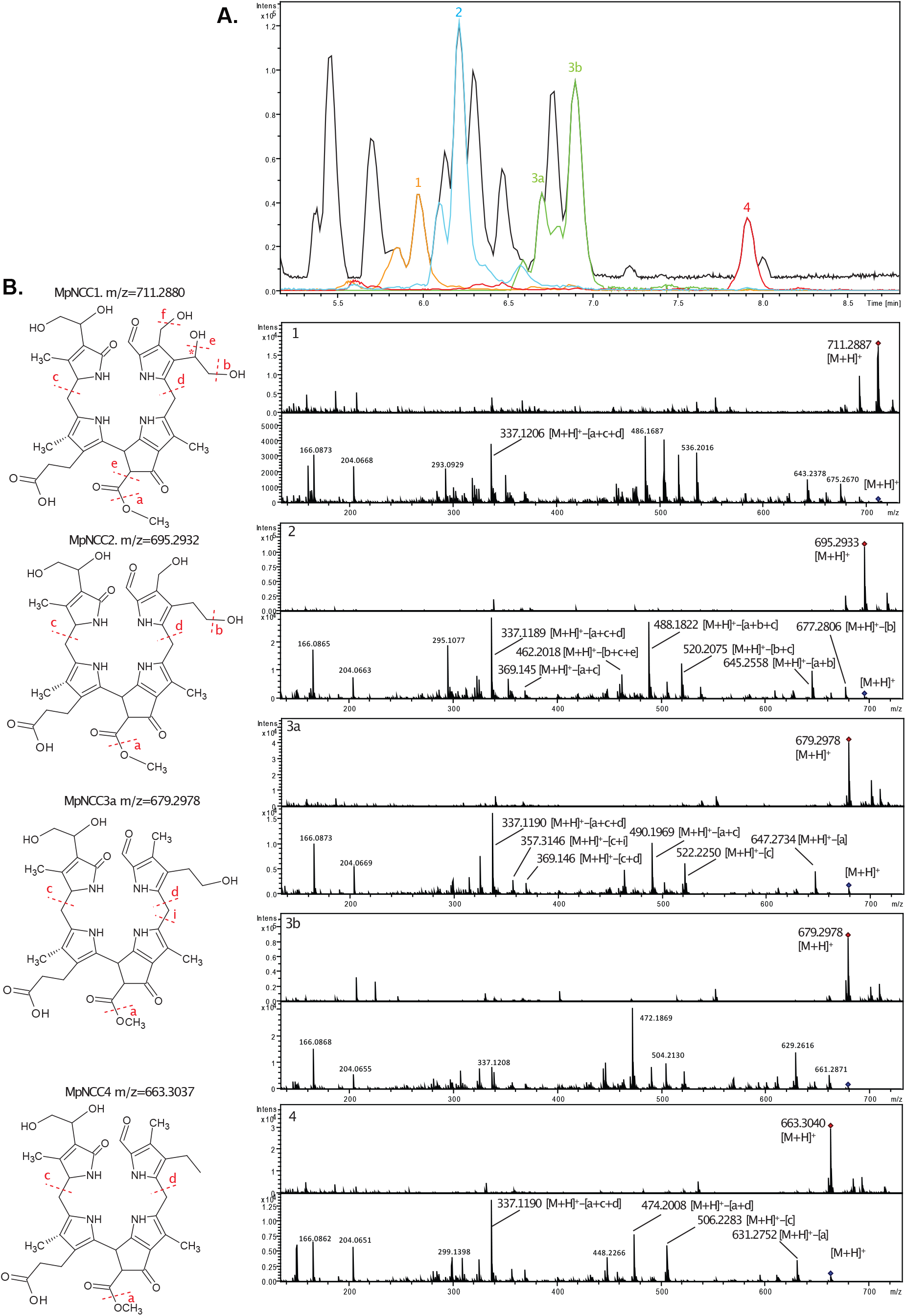
Phyllobilins detected in *Marchantia polymorpha. M. polymorpha* detached thalli were dark incubated for 10 days in the dark to induce chlorophyll degradation. **(A)** Base Peak chromatogram (in black) and extracted ion chromatogram of the four new phyllobilins detected by UPLC-MS/MS. **(B)** Putative structure models of all four phyllobilins detected and corresponding MS/MS fragmentation pattern. The red star in *Mp*NCC1 indicates a hypothetical position for the hydroxyl group that cannot be resolved based on the data available.

**Supplementary Figure 7.**
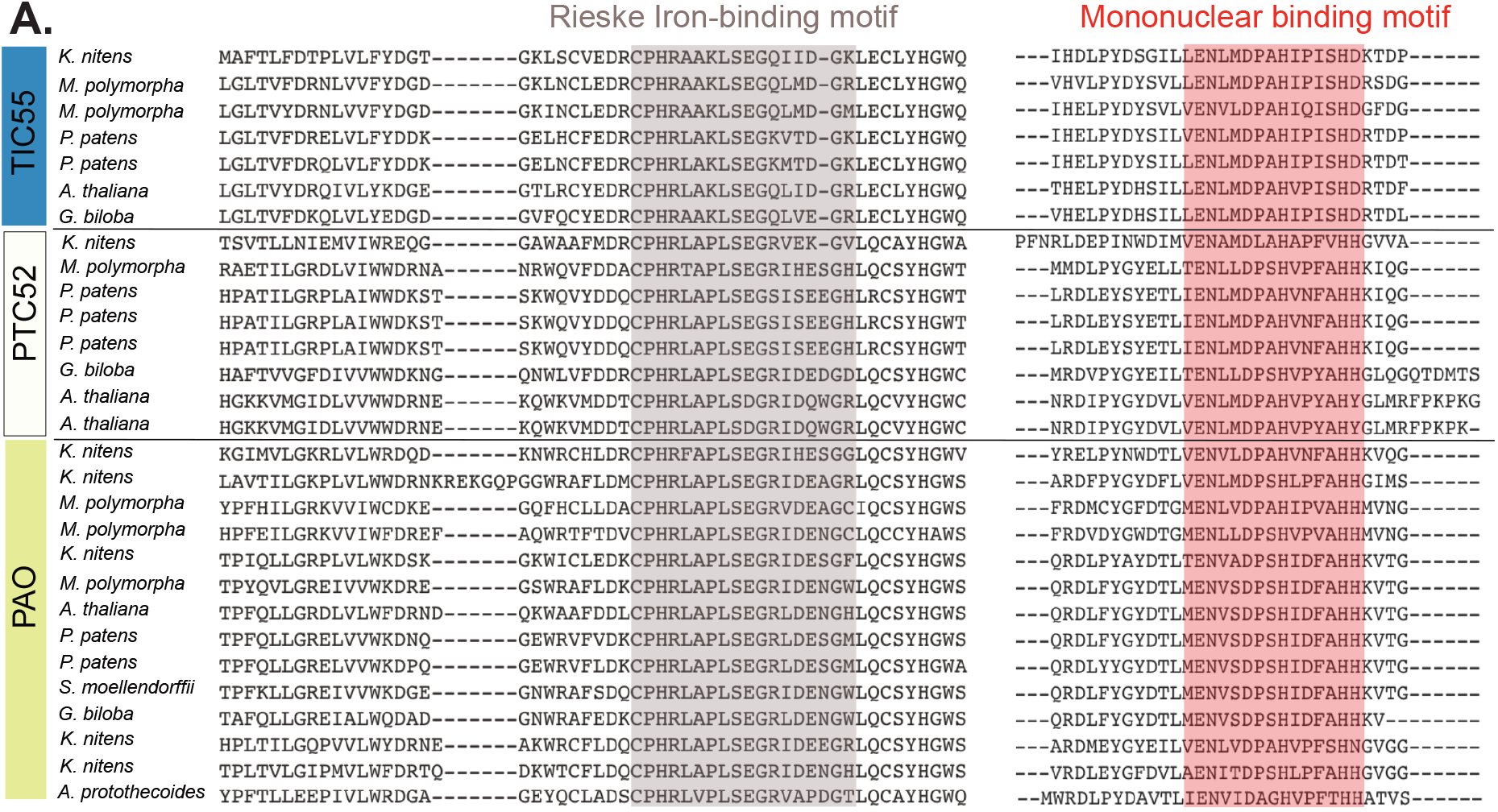
Alignment of Rieske and mononuclear iron domain of proteins from the PAO family. Alignment was performed using MAFFT and motif highlighted in colour.

